# A probability distribution over latent causes in the orbitofrontal cortex

**DOI:** 10.1101/041749

**Authors:** Stephanie C.Y. Chan, Yael Niv, Kenneth A. Norman

**Affiliations:** Princeton Neuroscience Institute, Princeton University, Princeton, NJ 08540

## Abstract

The orbitofrontal cortex (OFC) has been implicated in both the representation of “state”, in studies of reinforcement learning and decision making, and also in the representation of “schemas”, in studies of episodic memory. Both of these cognitive constructs require a similar inference about the underlying situation or “latent cause” that generates our observations at any given time. The statistically optimal solution to this inference problem is to use Bayes rule to compute a posterior probability distribution over latent causes. To test whether such a posterior probability distribution is represented in the OFC, we tasked human participants with inferring a probability distribution over four possible latent causes, based on their observations. Using fMRI pattern similarity analyses, we found that BOLD activity in OFC is best explained as representing the (log-transformed) posterior distribution over latent causes. Furthermore, this pattern explained OFC activity better than other task-relevant alternatives such as the most probable latent cause, the most recent observation, or the uncertainty over latent causes.

**SIGNIFICANCE STATEMENT:** Our world is governed by hidden (latent) causes that we cannot observe, but which generate the observations that we do see. A range of high-level cognitive processes require inference of a probability distribution (or “belief distribution”) over the possible latent causes that might be generating our current observations. This is true for reinforcement learning (where the latent cause comprises the true “state” of the task), and for episodic memory (where memories are believed to be organized by the inferred situation or “schema”). Using fMRI, we show that this belief distribution over latent causes is encoded in patterns of brain activity in the orbitofrontal cortex — an area that has been separately implicated in the representations of both states and schemas.

**CONFLICT OF INTEREST:** The authors declare no competing financial interests.

## INTRODUCTION

In recent years, cognitive neuroscientists studying reinforcement learning have recognized the importance of specifying representations of environmental “state” that capture the structure of the world in a predictive way (Gershman and Niv, 2010; Courville et al, 2006). At the same time, there has been renewed interest among cognitive neuroscientists in how memory encoding and retrieval are shaped by situation-specific prior knowledge (“schemas”, e.g. Tse et al, 2007). As work in this area progresses, it is important to clarify exactly what constitutes a schema and how schemas are formed.

Whether inferring the current “state” or the currently relevant “schema”, agents are making inferences about the hidden variables that underlie and generate our observations in the world. This inference can be concretely formulated in terms of Bayesian latent cause models (e.g., Gershman, Blei, and Niv, 2010). According to this framework, states and schemas can be viewed as hidden (latent) causes that give rise to observable events. For example, if you arrive late to a lecture, the situation (whether this is indeed the department colloquium or you have accidentally walked in on an undergraduate class) affects your observations about the average age of the audience, the proportion of audience members that are taking notes, the type of information being presented, and so on. To decide whether you are in the right place, you can use Bayesian inference to infer a belief distribution over the possible situations that might have generated the current observations, i.e. a posterior probability distribution over latent causes, *p(latent cause / observations)* (Figure 1A).

**Figure 1.**
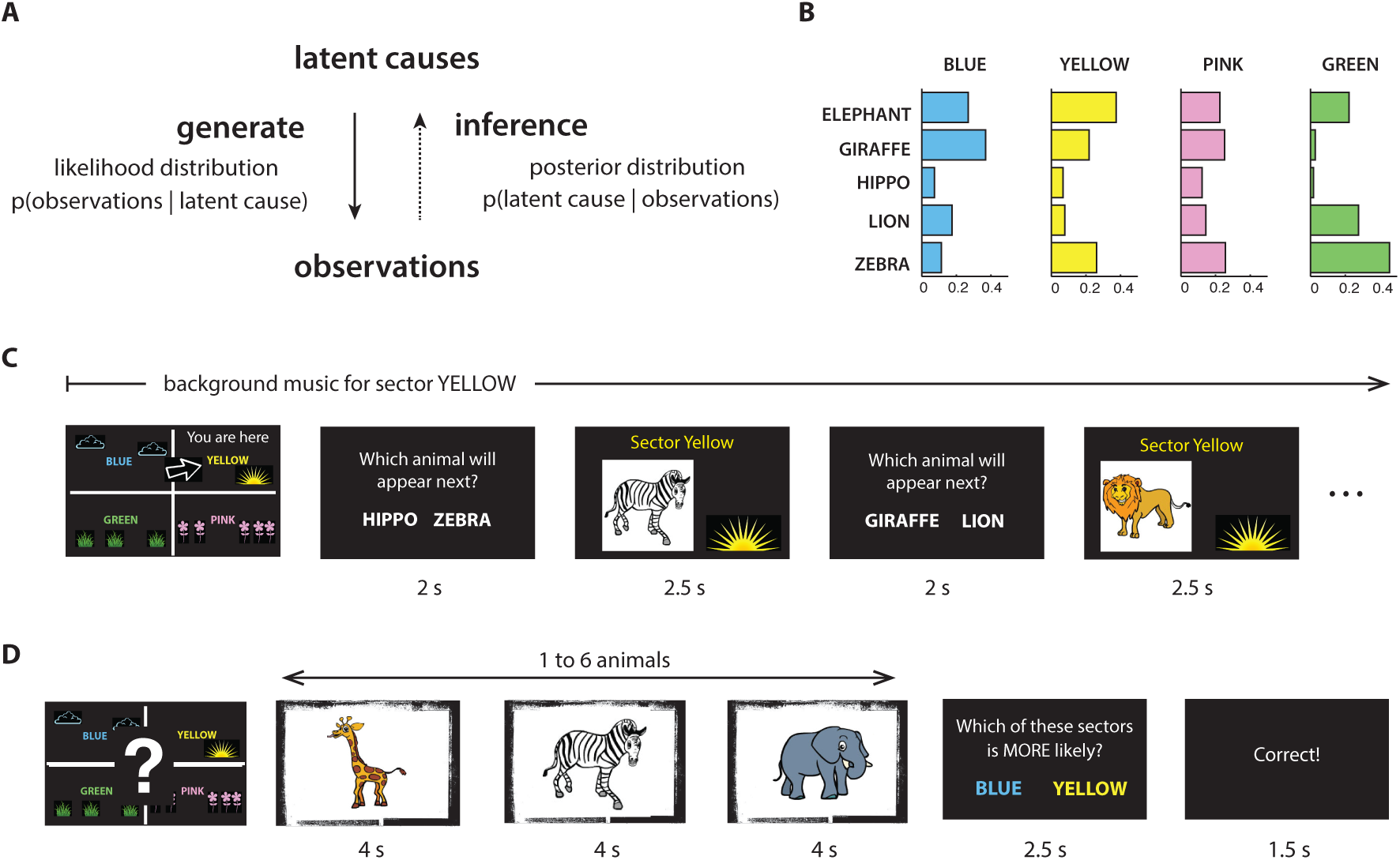
Task. ***A.** Schematic showing the relationship between latent causes and observations in the world. Inference about the posterior probability over latent causes involves inverting the generative model. **B.** Animal likelihood distributions P(animal | sector) (not shown directly to participants). Colors and animals were randomized across participants. **C.** An example of the first few trials of a tour through sector YELLOW. Each tour began with an image of the safari map, indicating the current sector and its location, and lasted 30-40 trials. Each trial began with a prompt asking the participant to guess which animal would appear next, followed by the appearance of an animal. A fixation cross was presented for 0.2-0.8 secs before each question and each animal presentation. The animals were pseudorandomly drawn from the likelihood distributions for the current sector. The sector’s music played in the background, until the start of the next tour. **D.** An example of a trial in the “photographs” task. Each trial began with an image of the safari map with a question mark at its center, indicating that the current sector was unknown. Next, a sequence of 1-6 animals appeared (pseudorandomly drawn from a single sector). Finally, participants were prompted to guess which of two sectors (randomly chosen) was more (or, on half the trials, less) probable. Participants received feedback on their responses. A fixation cross was presented for the last 0.5 secs of each animal presentation. [Thanks to sciencewithme.com for the animal illustrations.*]

We hypothesized, based on the similarity of the underlying computations, that the inference related to these two cognitive constructs (states and schemas) might be implemented using the same neural hardware. Indeed, there is one area of the brain that has separately been implicated in representing states (Wilson et al, 2014) and also schemas (Schlichting and Preston, 2015; Richards et al, 2014; Ghosh and Gilboa, 2014; Ranganath and Ritchey, 2012; van Kesteren et al, 2012; Tse et al, 2011) – the orbitofrontal cortex (OFC). Furthermore, previous univariate analyses in fMRI have implicated this region in encoding various summary statistical measures that are related to or are components of the posterior distribution, e.g. the posterior mean, likelihood of the current stimulus, and prior uncertainty (Ting et al, 2015; d’Acremont et al 2013; Vilares et al, 2012). However, these studies have not investigated representations of a full probability distribution.

Here, we used fMRI to investigate representation in OFC of posterior probability distributions over latent causes. In our experiment, we created a probabilistic environment in which participants were required to make inferences about the hidden causes that generated their observations. Participants viewed sequences of animal photographs, taken in one of four “sectors” in an animal reserve. They were tasked with judging the probability with which each sector generated the animal photographs, based on their previous experience observing animals in each sector. Using multivariate pattern similarity analyses of fMRI activity, we found that BOLD activity in the OFC was better explained by the posterior distribution over sectors (latent causes) than by a wide range of related signals, including the current stimulus, the most probable sector (the maximum a posteriori latent cause), or the uncertainty over latent causes (operationalized as the entropy of the posterior distribution). The present result advances our understanding of the function of the orbitofrontal cortex. It also unifies results from two different fields of cognitive neuroscience, inviting further investigation into the relationship between probabilistic inference, states, and schemas.

## MATERIALS AND METHODS

### Participants

32 participants (aged 18-34 years, 22 female) from the Princeton University community participated in exchange for monetary compensation ($20 per hour + up to $15 performance-related bonus). All participants were right-handed. Participants provided informed written consent. The study was approved by the Princeton University Institutional Review Board.

### Experimental design

#### The safari

Participants were told that they were going on a safari, visiting an animal reserve that was divided into 4 different sectors. Each sector was associated with a different color, background image, background music, and location on a 2 by 2 map (randomized across participants).

There were 5 different kinds of animals in the animal reserve. Every animal appeared in every sector, but with different likelihoods *P(animal | sector)*. The likelihoods (not shown directly to the participants) were chosen so that none of the sectors were strongly identified with a single animal, and so that none of the animals were strongly identified with a single sector (Figure 1B; colors and animals were randomly assigned across participants).

#### Procedure Overview

The experiment consisted of two parts. In the first part, participants “toured” through the animal reserve, in order to learn (through experience) the likelihoods *P(animal | sector)* for each animal and each sector. In the second part of the experiment, participants were shown sequences of “photographs” of animals that were taken in an unknown sector, and were asked to infer the posterior probabilities of different sectors given the animals shown in each sequence, *P(sector | animals shown)*.

For each participant, the experiment took place across two consecutive days (see Table 1).

**Table 1.** Tasks performed by participants on Day 1 and Day 2.

| Day 1 |  |  |
| --- | --- | --- |
| “Tours” task | 40 trials each tour | 2 tours through each sector, going clockwise around the map |
| “Tours” task | 20 trials each tour | 2 tours through each sector, sectors pseudorandomly ordered |
| Day 2 |  |  |
| “Tours” task | 30 trials each tour | 2 tours through each sector, sectors pseudorandomly ordered |
| “Tours” task | 10 trials each tour | 2 tours through each sector, sectors pseudorandomly ordered |
| “Photographs” task | 2 sessions x 20 trials each | outside of the MRI scanner |
| “Photographs” task | 4 sessions x 30 trials each | inside of the MRI scanner |

#### *“Tours”* task

In the “tours” task (Figure 1C), participants were instructed that they would “tour” through the animal reserve, one sector at a time, in order to learn the animal frequencies in each sector (the animal likelihoods). One animal appeared on each trial, pseudorandomly chosen according to the likelihoods for that sector. Before each animal appeared, participants were shown a prompt, asking them to make a prediction about which of two animals (one correct and one randomly chosen) would appear next. The alternate (incorrect) option was chosen with uniform probability from the four other animals. To distinguish between the animals in the question prompt (which were not representative of the sector’s likelihood distribution) and the animals that were actually drawn from the sector’s likelihood distribution, the question prompts were shown as text while the animals drawn from the safari sector were shown as pictures.

In order for the sectors to form rich contexts, each sector was associated with a different color, background image, background music, and location on a 2 by 2 map (randomized across participants). Before the first trial of a tour through a sector, participants were shown the sector’s location on the map. Also, for the duration of a tour through a sector, animals were displayed on the sector’s color-matched backdrop image, and the music associated with that sector was played in the background.

#### *“Photographs”* task

On each trial of the “photographs” task (Figure 1D), participants were shown a sequence of animal “photographs”, without being told which sector the photographs were taken from. At the end of the sequence, participants were prompted to indicate which of two sectors (randomly chosen) was more (or less) probable. The two sector options for each question were chosen uniformly from the four sectors of the safari (and did not necessarily include the most or least likely sector). So, to perform well on the task, participants had to maintain a full posterior distribution over all four sectors (as opposed to estimating only the most probable sector, for instance).

Participants received 10 cents for every correctly answered question, and they received feedback on every trial. So that more probable sectors were not consistently associated with higher monetary value, we asked which of the two sectors was *more* probable on half of the trials, and which was *less* probable on the other half of the trials. To eliminate confounds with motor plan, the positions of the two response options were pseudorandomly assigned between left and right.

To encourage participants to update their inference of the sector probabilities after every animal presentation (as opposed to waiting until the time of the question to integrate over the animals observed), we varied the length of the sequences between 1 and 6 animals (so that the appearance of the question prompt was unpredictable), and participants were only allowed 2.5 seconds to give a response after the appearance of the question.

The posterior probability of each sector *P(sector | animals seen)* can be straightforwardly computed from the animal likelihoods, using Bayes rule (all sectors were equally likely a priori):

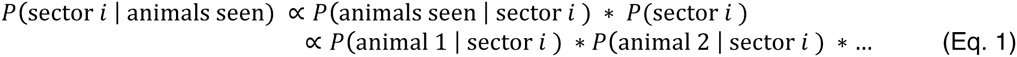

Feedback for the responses was generated based on these posterior probabilities. Due to a bug in the code that was undetected during data collection, the feedback was incorrectly generated for some of the trials containing only one animal presentation (this affected approximately 10% of the trials). In our fMRI analyses, to account for learning from the incorrect feedback, we used each participant’s estimates of the likelihoods (collected at the end of the experiment) instead of the real likelihoods, and we also performed trial-by-trial behavioral model-fitting to model learning from feedback (see next section).

Participants first performed 2 sessions (20 trials each) of the “photographs” task outside the MR scanner, to familiarize themselves with the task. They then performed 4 sessions (30 trials, approximately 11 minutes per session) inside the scanner.

### Behavioral model-fitting

To model participants’ posterior inference on the “photographs” task, as well as any learning from feedback, we performed trial-by-trial model-fitting of participants’ responses. We tested several classes of models:

#### Bayesian_nolearning

This model assumed that participants were correctly computing the posterior distribution over sectors *P(sector | animals seen)* using Bayesian inference (as in Eq. 1). To obtain the model-derived likelihood of each behavioral response (and to capture stochasticity in participants’ behavior), we used a softmax on the posterior probabilities of the two options in each question prompt.

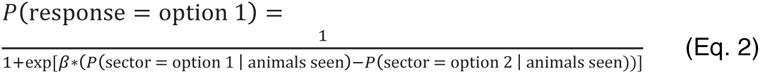

where β is an inverse temperature parameter (**(3**= 0 implies equal likelihood for both options).

#### additive

In this model, instead of correctly multiplying the animal likelihoods together to obtain the posterior distribution over sectors (as in Eq. 1), we assumed that participants *added* the likelihoods together to obtain an “additive posterior” (normalized to sum to 1).

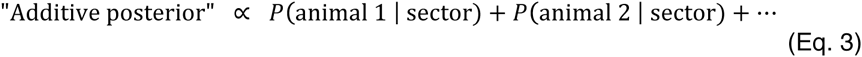

While statistically suboptimal, we might expect this from a simple associative mechanism that brings the sectors to mind in proportion to their association strength with the animals seen. Again, to determine response probabilities, we applied a softmax operator to the additive “posterior” probabilities for the two options in each question prompt.

#### most/least voter

These models assumed that participants were only paying attention to the most common (and/or least common) animals in each sector, a similar strategy having been previously observed in a similar task (Gluck et al, 2002). During the trials, each animal appearance "voted" for (or against) the sectors in which it was the most common (or least common). To obtain the model-derived likelihood of each behavioral response, we used a softmax on the final tally at the end of each sequence.

We tested several variants of this model, e.g. tallying only the positive votes, and/or allowing an animal to “vote” for (or against) a sector if it was one of the *two* most (or least) common animals in that sector. The magnitude of the positive and negative votes were either allowed to be two separate free parameters, or constrained to be equal to each other. Because the magnitude of the vote already served as a scaling parameter for the input to the softmax operator, the inverse temperature of the softmax was kept constant at 1.

#### Bayesian feedbackRL

These models were designed to account for learning from feedback during the “Photographs” task (including the incorrectly generated feedback). Here we assumed a reinforcement learning process, in which participants adjusted their internal estimates of the animal likelihoods after feedback about the two sectors in the question. These likelihoods were then used to compute the posterior distribution over sectors via Bayes rule.

For the sector that feedback indicated to be more probable, likelihoods were adjusted upwards for all animals that were seen on that trial. For the sector that was indicated to be less probable, likelihoods were adjusted downwards for all animals seen on the trial (see Figure 2 for an example).

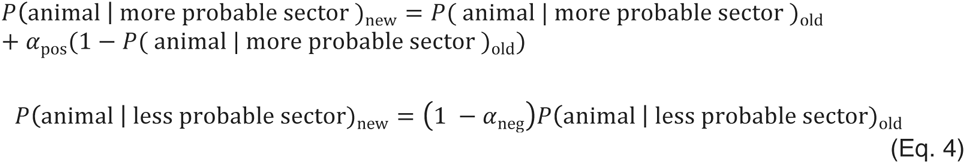

**Figure 2.**
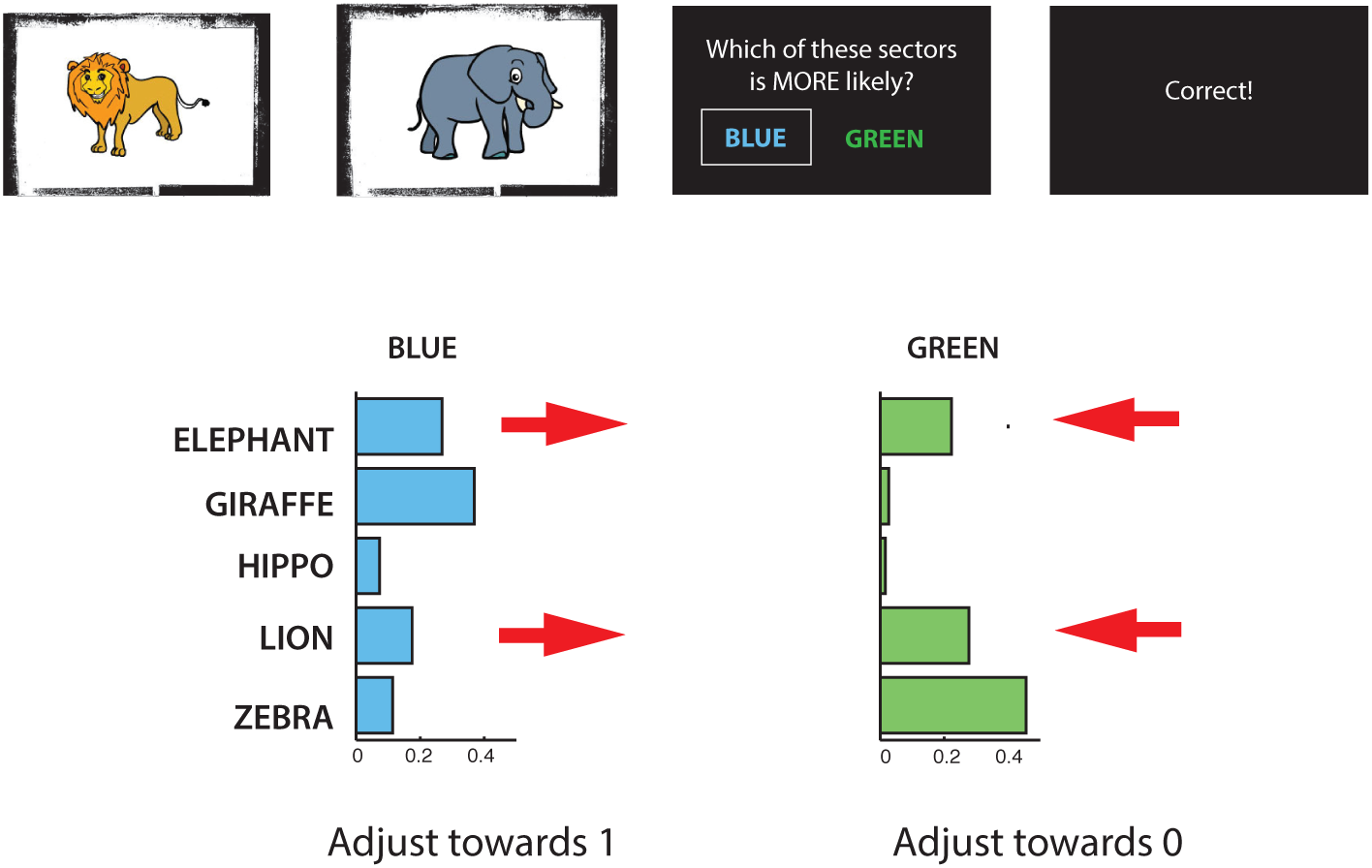
FeedbackRL model. *An illustration of learning from feedback in the Bayesian_feedbackRL model, for a single trial (not real data). In this example trial, the participant saw a lion and an elephant, and was asked about sector BLUE and sector GREEN. The feedback indicated that sector BLUE was more probable. As a result, the likelihoods P(BLUE | elephant) and P(BLUE | lion) are adjusted towards 1 with learning rate α_pos_, and the likelihoods P(GREEN | elephant) and P(GREEN | lion) are adjusted towards 0 with learning rate α*_*neg*_.

Estimates of the likelihoods were renormalized after each adjustment. The learning rates α_pos_ and α_neg_ were either allowed to be two separate free parameters, or they were constrained to be equal.

For the initialization of the likelihoods, we tested two versions of the model: initialization at the true animal likelihoods, or initialization according to the participants’ subjective estimates of the likelihoods (collected at the end of the experiment, see below).

Finally, the likelihoods were used to compute the posterior distribution over sectors via Bayes rule. Thus, posterior inference in the FeedbackRL model also used Bayes rule – the only difference from the “Bayesian_nolearning” model above is that the likelihoods (which enter into the posterior inference computation from Eq. 1) were adjusted on each trial according to feedback.

We tested several additional variants of this model. In one variant, participants only adjusted their likelihoods in response to “You are incorrect” feedback (instead of in response to all feedback). In another variant of the model, we scaled the learning rates separately for each animal according to how much that animal contributed to the final posterior distribution:

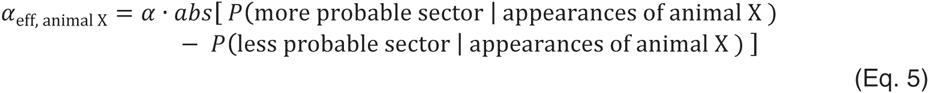

In this variant, animals appearing multiple times in a trial would have higher effective learning rates, having contributed more to the final decision.

In a post-experiment questionnaire, we asked participants to provide their estimates for the animal likelihoods in each sector. For each of the models above, we tested versions using (a) the *actual* animal likelihoods, and (b) *subjective estimates* of the animal likelihoods. For the few participants who provided likelihood estimates that did not sum to 1, we normalized the estimates. To avoid taking logarithms of 0, we converted estimated likelihoods of 0 to 0.01 (and renormalized).

For each of the models, we also tested versions in which the earlier and/or later animals in each sequence were given extra weight. To model these primacy/recency effects, we fit a power law function for each participant to give more weight to the earlier and/or later animals in each sequence (e.g. 1^*w*^, 2^*w*^, … for animal 1, animal 2, …). The likelihoods were exponentiated by this weighting and renormalized. If modeling both recency and primacy, the weightings for each were summed. We tested versions in which the recency and primacy free parameters *w* were either allowed to be two free parameters, or they were constrained to be equal.

Free parameters for each model were fit to each participant’s behavioral data separately, using Matlab’s “fmincon” function, with at least ten random initializations for each model and each participant. The best-fitting parameters (the maximum likelihood estimates) were used to evaluate, for each participant and each model, the (geometric) mean likelihood per trial (i.e., the exponentiated log likelihood per trial, without any penalization for number of parameters), the Akaike information criterion (AIC), and the Bayesian Information Criterion (BIC), in order to compare the models and determine which best accounted for participants’ behavior.

#### fMRI acquisition and pre-processing

Functional brain images were acquired using a 3T MRI scanner (Siemens, Skyra) and preprocessed using FSL (http://fsl.fmrib.ox.ac.uk/fsl/). An echoplanar imaging sequence was used to acquire 36 slices (3mm thickness with 1mm gap, repetition time (TR) = 2s, echo time (TE) = 27ms, flip angle = 71º). To increase signal in the OFC, slices were angled approximately 30 degrees from the axial plane towards a coronal orientation (Deichmann et al, 2003). For each participant, there were 4 scanning runs in total (approximately 11 minutes each). The functional images were spatially filtered using a Gaussian kernel (full width at half maximum of 5mm), and temporally filtered using a low-pass cutoff of 0.0077Hz. We performed motion correction using a six-parameter rigid body transformation to co-register functional scans, and then registered the functional scans to an anatomical scan using a 6-parameter affine transformation.

The motion regressors (and their derivatives) were residualized out from the functional images, as were the mean timecourses for cerebrospinal fluid and white matter (segmentation was performed using FSL’s “FAST” function), and also the mean timecourse for blood vessels (estimated by taking voxels with the top 1% in standard deviation across time). Then, the functional images were z-scored over time. All analyses were performed for each participant in participant space, and then spatially normalized by warping each participant’s anatomical image to MNI space using a 12-parameter affine transformation.

#### Region of interest – Suborbital sulcus

Our region of interest (ROI) was determined as the intersection of two sets of brain areas. The first set of areas, the orbitofrontal cortex, has been postulated to be involved in the representation of “state”, due to evidence from studies of human and animal reinforcement learning (Wilson et al, 2014). The second set of areas, sometimes referred to as the “posterior medial network”, has been postulated to be involved in the computation and representation of “schemas” or “context” (Ranganath and Ritchey, 2012), as the set of areas with high connectivity with parahippocampal cortex (PHC). The intersection of these sets of areas is the suborbital sulcus, a medial subregion of the orbitofrontal cortex (Figure 6A). Using Freesurfer (Destrieux et al, 2010), the ROI was drawn as the anatomically parcellated cortical region centered on the voxel with maximal resting-state functional connectivity to PHC (Libby et al, 2012).

**Figure 3.**
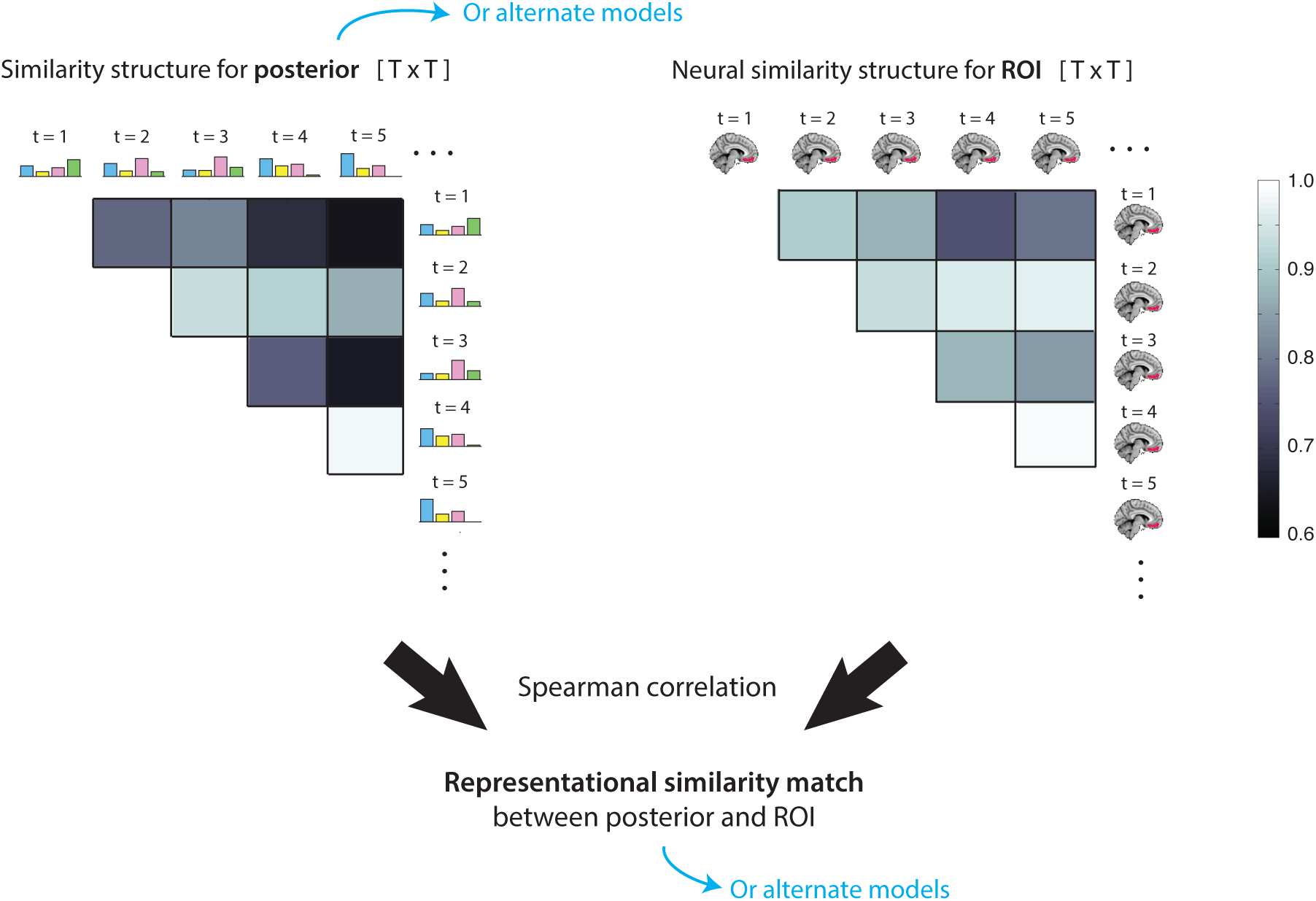
Representational similarity analysis. *An illustration of the representational similarity analysis (not real data). We first computed the similarity structure for the posterior distribution (or any alternative model; see Table 3) by computing the normalized correlation of the posterior at every timepoint with every other timepoint. We also computed the neural similarity structure for our region of interest (or for each searchlight in the whole-brain analysis), by computing the normalized correlation between patterns of activity at every timepoint with every other timepoint. To evaluate the representational similarity match between the neural data and the model, we then computed the Spearman correlation between the two matrices (using only the upper triangle of each matrix, excluding the diagonal*).

**Figure 4.**
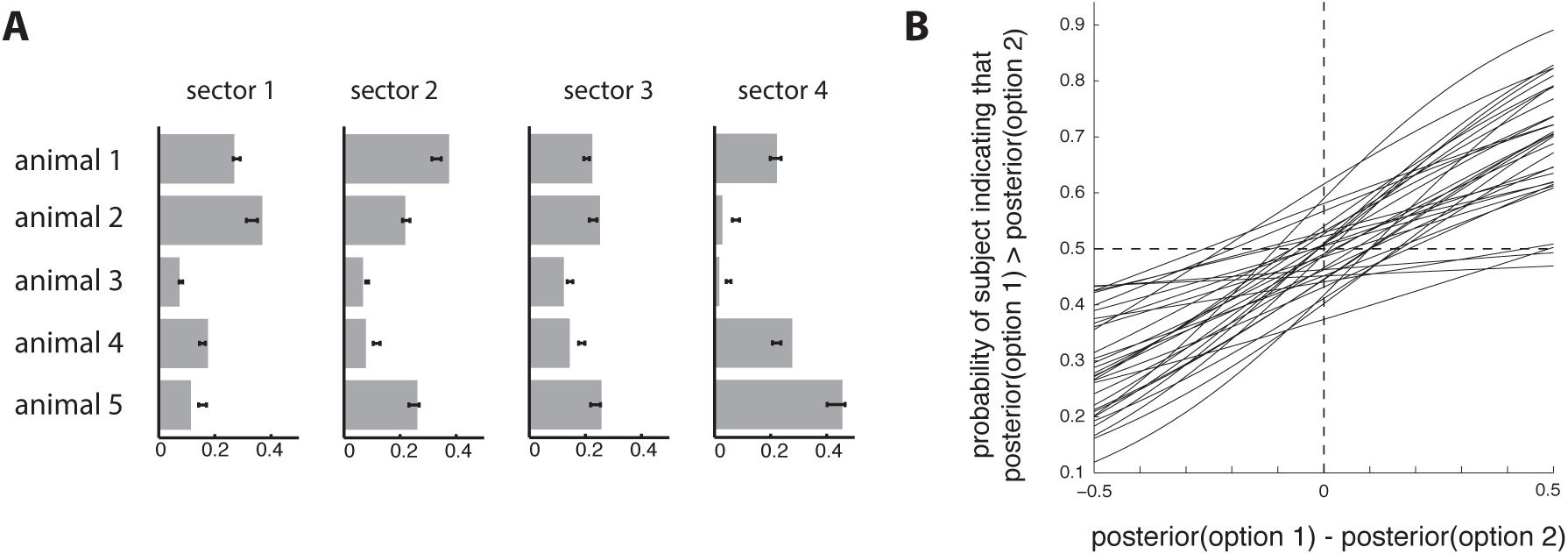
Behavioral performance. ***A.** Participants’ subjective estimates of the animal likelihoods P(animal | sector), for each animal and each sector, collected in a post-experiment questionnaire. Gray bars indicate the true likelihoods, black intervals indicate the mean estimates ± SEM. **B.** Logistic regression on participants’ responses during the fMRI scan sessions suggests that participants learned and utilized the full posterior distributions (each line shows logistic regression for one participant). The x-axis indicates the difference in posterior probability between the first and second options in the question. The y-axis indicates the probability that a participant would indicate that the first option has higher posterior probability than the second option. Mean regression parameters across participants: slope = 1.8 ± 0.040, intercept = -−0.04 ± 0.15*.

**Figure 5.**
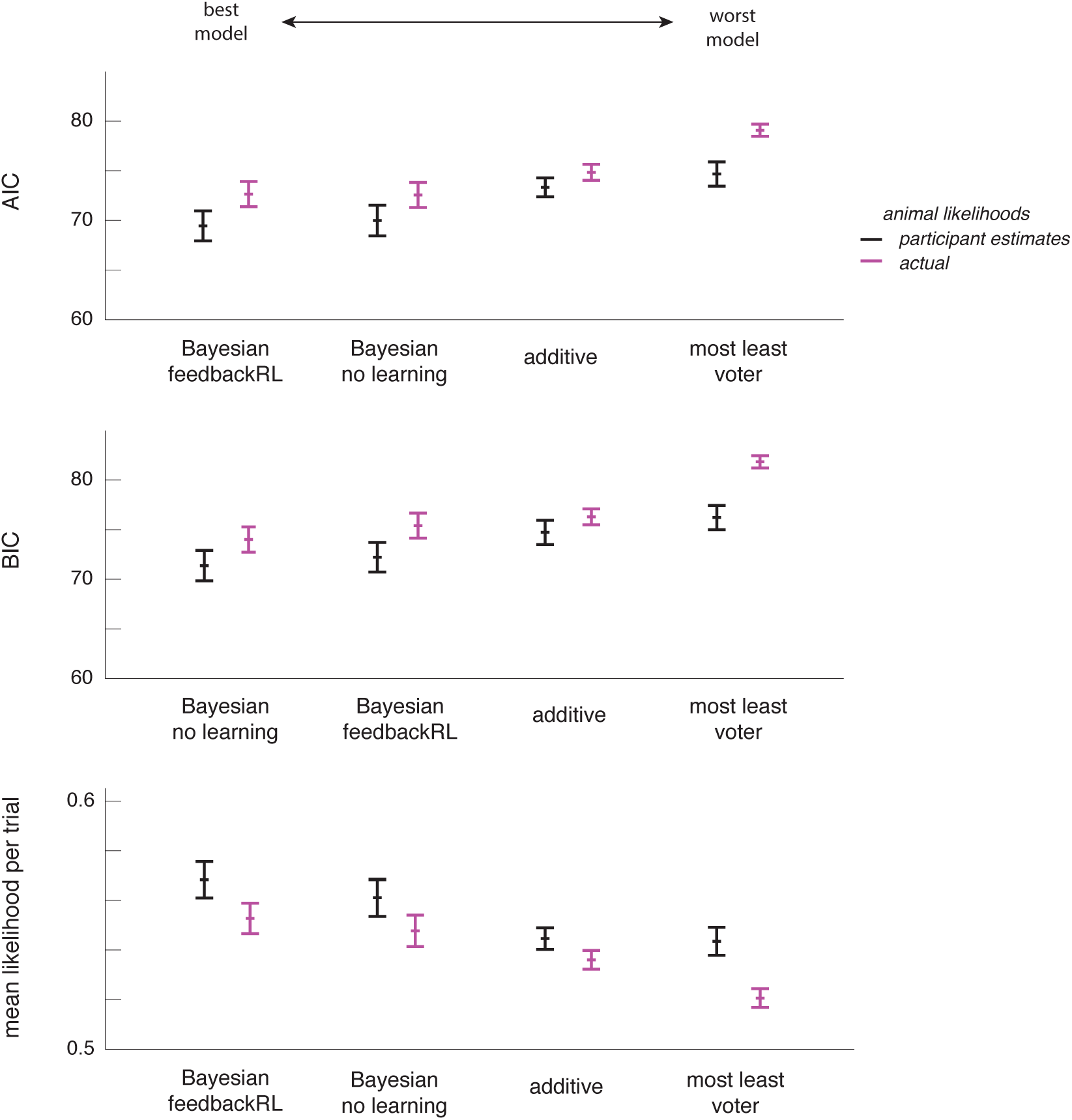
Behavioral model-fitting. *Akaike information criterion (AIC), Bayes information criterion (BIC), and (geometric) mean likelihood per trial (i.e. the exponentiated mean log likelihood per trial, without penalization for number of parameters) for the best-fitting model in each class (mean ± SEM across participants) suggest that the Bayesian models explained the behavioral data best. Note that better model fits are indicated by low AIC and BIC scores, but high mean likelihood. Results are shown for model fits using the participant estimates of the likelihoods or using the actual (true) likelihoods*.

**Figure 6.**
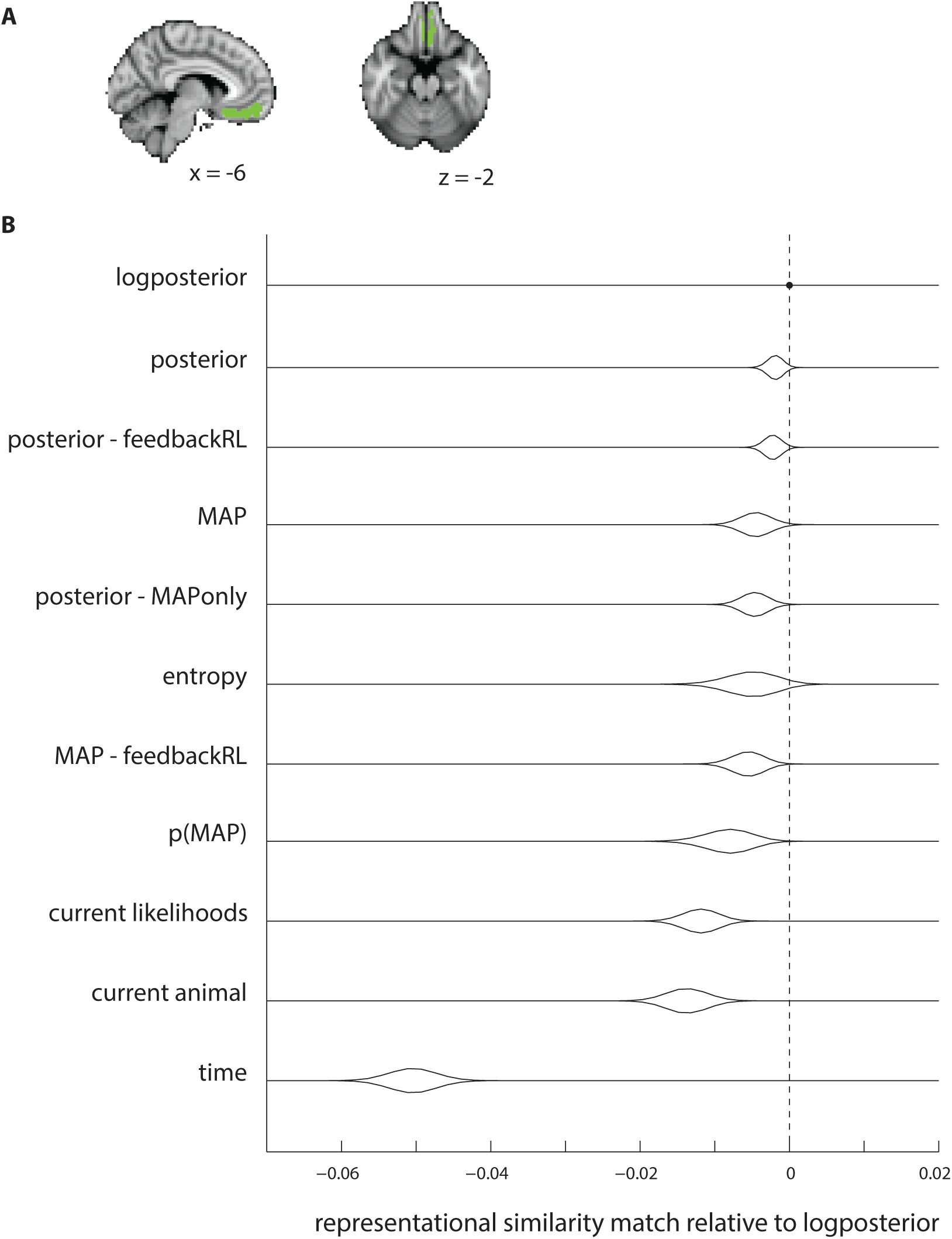
Representational similarity match for each model in the ROI. ***A.** Region of interest – the suborbital cortex, a medial subregion of the orbitofrontal cortex (OFC). See Methods for a description of how the region was defined. **B.** Representational similarity match for each of the models tested (Table 3), relative to the best model for the data (the logposterior), ordered by mean representational similarity match. The logposterior model showed the highest mean representational similarity match. The plots show bootstrap distributions on the within-participant differences, for each of the models compared with the logposterior. For all of the alternative models tested, 95% or more of our bootstrap samples showed a better match for the logposterior than for the alternative model*.

#### Representational similarity analysis

If the suborbital region of interest (ROI) contains a multivariate representation of the posterior distribution over latent causes, then patterns of neural activity in this area should be more similar for pairs of timepoints at which the posterior distribution was similar, and they should be dissimilar for pairs of timepoints at which the posterior distribution was dissimilar. Therefore, to test whether multivariate patterns of activity in the ROI might be representing the posterior distribution over sectors, we performed a representational similarity analysis (RSA; Kriegeskorte et al, 2008).

We first computed the similarity of the posterior distribution over sectors for every pair of timepoints during which we expected the posterior distribution to be updated (i.e. at the times of the animal appearances). This provided us with the *similarity matrix for the posterior.* We also computed the similarity of the neural pattern in the ROI for every pair of timepoints—the *similarity matrix for the ROI*. Then we computed the Spearman rank correlation of these two matrices (taking only the upper triangle and excluding the diagonal). We denote this Spearman correlation as the *similarity match between the posterior and the ROI* (Figure 3). We expected the similarity match to be positive, i.e. that the neural patterns in the ROI should be more similar for pairs of timepoints at which the posterior distribution over sectors was more similar.

We also computed the similarity match for the ROI with other signals, to compare with the similarity match between the ROI and the posterior distribution over latent causes. This is important because the similarity structure for the ROI could potentially be correlated with the similarity structure of the posterior distributions for reasons other than the fact that the posterior distribution is represented in this area. For example, the posterior distribution is, on average, more similar for pairs of timepoints at which the same animal is presented—if the suborbital ROI represents the animal currently presented, we would also find a positive similarity match between the ROI and the posterior distribution. We therefore compared the similarity match between the ROI and each alternate model, to determine the model that best explained the similarity structure of the neural data.

The set of alternate models used for this comparison included the log-transformed posterior distribution (since many signals in the brain are known to be represented in log space; e.g. Yang and Shadlen, 2010; Gibbon, 1977; Longo and Lourenco, 2007), the current stimulus, the maximum a posteriori (MAP) sector (most probable sector), the entropy of the posterior distribution (a proxy for overall uncertainty), the probability of the maximum a posteriori sector (a proxy for confidence, acting approximately as the converse of the entropy), and temporal distance between measurements (because fMRI pattern similarity is known to vary as a function of the temporal distance between measurements). We also included models of the posterior and MAP that were instead derived using the Bayesian_feedbackRL model (given that this was the best inference model after Bayesian_nolearning, as determined from behavioral model-fitting, described above). See Table 3 for a full list of models tested.

**Table 3.**
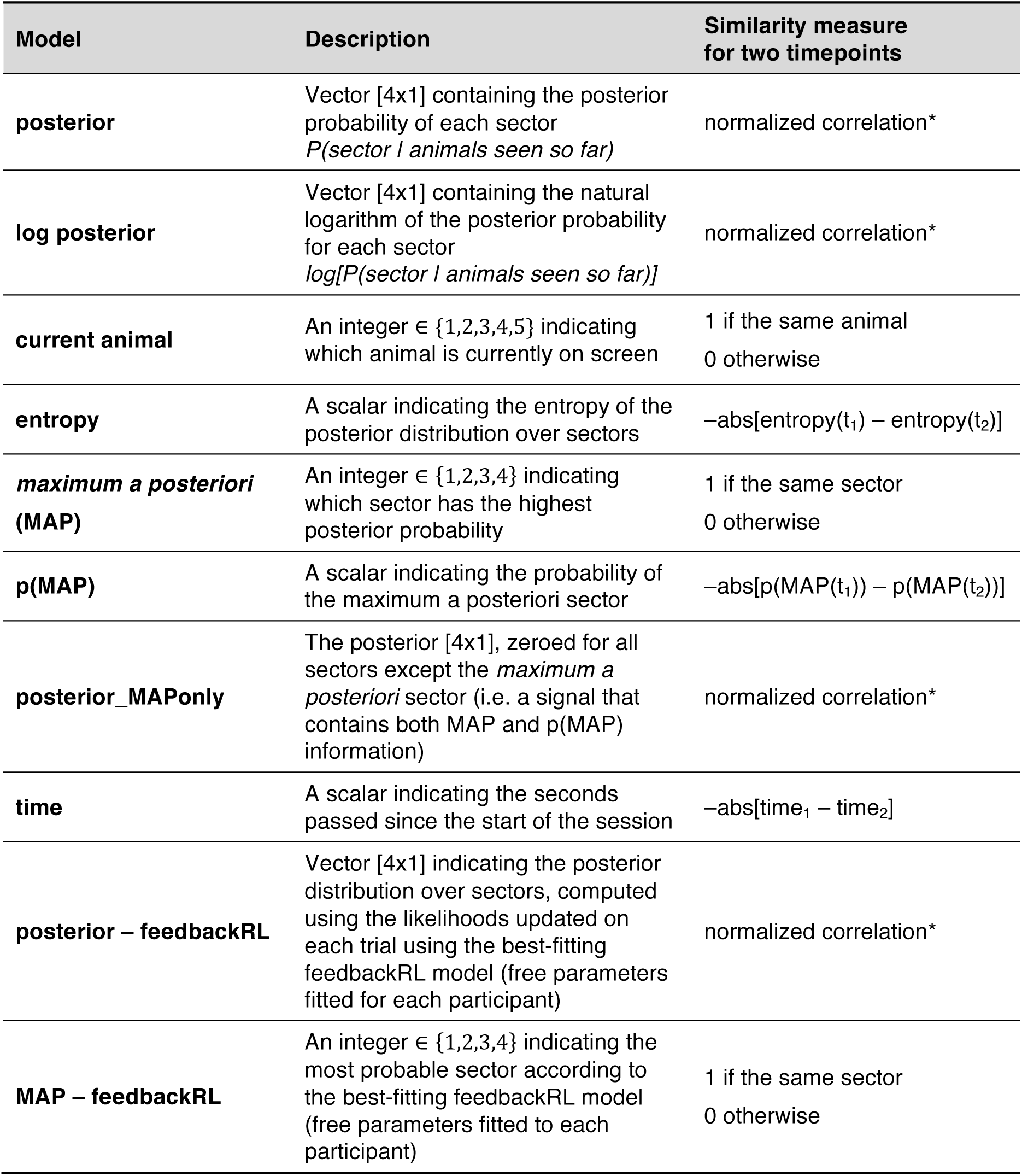
Models used in the representational similarity analysis, and the similarity measure used to derive the similarity matrix. *The normalized correlation of vectors **x** and **y** is **x** • **y**/(||**x**|| * ||**y**||), and is equivalent to the cosine of the angle between the two vectors. It behaves differently than the more commonly used Pearson correlation; for example, the posterior distributions [0.24 0.25 0.25 0.26] and [0.26 0.25 0.25 0.24] have Pearson correlation of −1 but normalized correlation of 0.9994. We used normalized correlation because this measure accords better with intuition regarding the similarity of posterior distributions and quantities derived from posterior distributions; however, similar results were observed when using Pearson correlations instead.

To investigate the specificity of the result to our region of interest, we also performed a whole-brain “searchlight” analysis, using 25-voxel spherical searchlights. As with the region of interest, we computed the similarity of the neural patterns in each searchlight, to obtain the *neural similarity matrix for the searchlight*. We then computed the Spearman correlation of the similarity matrix for each searchlight with each of our models. The analysis was repeated for a searchlight centered on every voxel in the brain.

For both the ROI and searchlight analyses, the neural pattern for each animal appearance was averaged over the two TRs during which the animal appeared on the screen (after correcting for the hemodynamic lag with a 4 second shift). Similarity for neural patterns was computed using normalized correlation, to accord with the similarity measure used for the posterior-based models (similar results are obtained when using Pearson correlation instead). Searchlight results are displayed on an inflated brain, using the AFNI SUMA surface mapper (http://afni.nimh.nih.gov/afni/suma).

#### Statistics and confidence intervals

Unless stated otherwise, all statistics were computed using random-effects bootstrap distributions on the mean by resampling participants with replacement (Efron & Tibshirani, 1986). All confidence intervals in the text are given as standard error of the mean.

To test the reliability of searchlight results across participants, we used the “randomise” function in FSL (http://fsl.fmrib.ox.ac.uk/fsl/fslwiki/randomise) to perform permutation tests and generate a null distribution of cluster masses for multiple comparisons correction (using FSL’s “threshold-free cluster enhancement”, *P* < 0.05 two tailed).

## RESULTS

### Participants learned the animal likelihoods in the “Tours”

We evaluated participants’ final learning of the likelihood of each animal in each sector using performance from the last set of tours on the last day. In those tours, the participants chose the more likely animal 73 ± 3% of the time. Note that even if participants had perfect knowledge of the animal likelihoods, we would not expect participants to choose the more likely animal 100% of the time, due to probability matching, the well-documented behavior in which humans and animals match their choice probabilities to the probability of each option being correct, rather than choosing the most likely option every time (e.g. Vulkan, 2000; Erev and Barron, 2000). With perfect knowledge of the animal likelihoods and a probability matching policy, participants would be expected to choose the more likely animal only 69% of the time.

In a post-experiment questionnaire, we asked participants to estimate the animal likelihoods *P(animal | sector)* for every animal and every sector. These estimates were close to the true likelihoods, on average (Figure 4A). The mean KL-divergence of the estimated likelihoods from the real likelihoods was 0.13 ± 0.015. As discussed below, we used these participant-estimated likelihoods in our neural analyses, in lieu of the correct likelihoods.

### Performance on “Photographs” task suggested maintenance of posterior distributions over sectors

During the fMRI scan sessions, participants correctly chose the more (or less) probable sector 67 ± 1% of the time, which is significantly above chance (t-test *p* < 1e-12). Moreover, logistic regression on participants’ responses showed that, the larger the difference in posterior probability between the correct and incorrect options, the more likely participants were to choose the correct answer (Figure 4B). Again, as in the Tours task, we expected stochasticity in participants’ behavior due to probability matching. With perfect probability matching and perfect inference of the sector posteriors, we would expect participants to choose the correct option 73% of the time.

Note that the two sector options in each question were chosen at random, and therefore required participants to discriminate between posterior probabilities for any possible pair of sectors. Interestingly, participants performed similarly well whether or not questions included the maximum a posteriori (MAP; most probable) sector (accuracy 69 ± 2% for questions including the MAP, 66 ± 1% for questions not including the MAP; not significantly different). This result further indicates that participants were tracking the full posterior distribution, and not just the MAP sector.

### Trial-by-trial behavioral model-fitting suggested that participants were approximately Bayesian

The relative performance of the behavioral models is shown in Figure 5, and the mean parameter fits are shown in Table 2. For model comparison, we used the best-performing version from each class of models (these settings described in Table 2).

**Table 2.** Free parameters and parameter fits, for the best-fitting model for each class. *The best-fitting models for all classes did not model recency or primacy biases, and used each participant’s subjective estimates of the animal likelihoods rather than the actual likelihoods. For model classes that had additional variants, the best-fitting settings are described in parentheses*.

| Model | Free parameters | Mean ± SE | Range |
| --- | --- | --- | --- |
| **Bayesian_nolearning** | $\beta$ - softmax inverse temperature | $4.04 \pm 2.22$ | $[0, \infty]$ |
| **additive** | $\beta$ - softmax inverse temperature | $7.04 \pm 3.46$ | $[0, \infty]$ |
| **mostleast_voter**<br>(voting for or against the sectors in which an animal was the most or least common) | $v_{\text{pos}}$ - size of positive vote<br>$v_{\text{neg}}$ - size of negative vote | $1.69 \pm 3.39$<br>$0.754 \pm 1.39$ | $[0, \infty]$<br>$[0, \infty]$ |
| **Bayesian_feedbackRL**<br>(learning from all feedback, no scaling of learning rates, and $\alpha_{\text{pos}} = \alpha_{\text{neg}}$) | $\alpha$ - learning rate<br>$\beta$ - softmax inverse temperature | $0.0515 \pm 0.161$<br>$4.82 \pm 2.52$ | $[0, 1]$<br>$[0, \infty]$ |

The two Bayesian models (with and without feedbackRL) performed best, explaining the data about equally well. Overall, the model with feedbackRL was the best model according to AIC, but the Bayesian model without learning was the best model according to BIC, which penalizes more strongly for extra parameters.

The additive model performed worse than the Bayesian models, indicating that participants were accumulating evidence multiplicatively, in accordance with the optimal strategy (Eq. 1). None of the heuristic inference models that we tested (the most-least voter class of models) could successfully outperform the Bayesian models. Nor did we identify any significant effect of recency or primacy (any small improvements in the model likelihoods were not justified by the increased number of parameters). We therefore concluded that participants were Bayesian or near-Bayesian in their inference.

As shown in Figure 5, using the participants’ subjective estimates of the animal likelihoods (from the post-experiment questionnaire) provided a better fit for all models, as compared to using the real animal likelihoods. This may be surprising for the feedbackRL model, given that the participant estimates were elicited at the end of the experiment, but were used in the model to initialize estimates of the likelihoods. However, the low learning rates (see Table 2 for average fit learning rates; also, 19% of participants had fitted learning rates of 0) suggest that changes in the likelihoods throughout the experiment were small relative to the differences between the real and estimated likelihoods. The low learning rates also explain why the feedbackRL model fit the data similarly well to a Bayesian model that did not allow for changes of the likelihood during the task – the models are nested (identical for learning rates of zero) and similar for low learning rates.

### Representational similarity analysis suggests that suborbital sulcus contains a representation of the (log) posterior distribution over latent causes

Figure 6B shows the representational similarity match of the suborbital sulcus with each of the models, relative to the representational similarity match with the best model – the logposterior. For all of the alternative models tested, 95% or more of our bootstrap samples showed better representational similarity match for the logposterior than for the alternative model.

Because the posterior distribution tends to be more similar for neighboring timepoints compared with more distant timepoints, and that might also be the case for neural patterns, we took special care to verify that the logposterior model was superior to the alternative (control) time model. This was indeed the case. Moreover, we found that the temporal model displayed negative representational similarity match with the neural patterns, because BOLD patterns for neighboring timepoints tended to be anti-correlated. This result was not dependent on our linear model for temporal distances—because we used Spearman’s rank correlation to compute representational similarity match, the negative similarity match result would be observed for any other model of temporal distance that falls off monotonically (e.g. an exponential model). Therefore, since the posterior distribution showed positive similarity match while the temporal model showed negative similarity match, we can conclude that any positive correlations between the similarity matrices for the posterior distribution and time cannot be responsible for the representational similarity result for the posterior distribution.

Searchlight results for the representational similarity analysis are shown in Figure 7. The orbitofrontal and ventromedial prefrontal cortex showed significantly greater representational similarity match for the logposterior model compared to every other model (p < 0.05 corrected, for every comparison), except for the entropy and the posterior models. It also showed greater representational similarity match for the logposterior than entropy using a more liberal threshold of p < 0.05 uncorrected.

**Figure 7.**
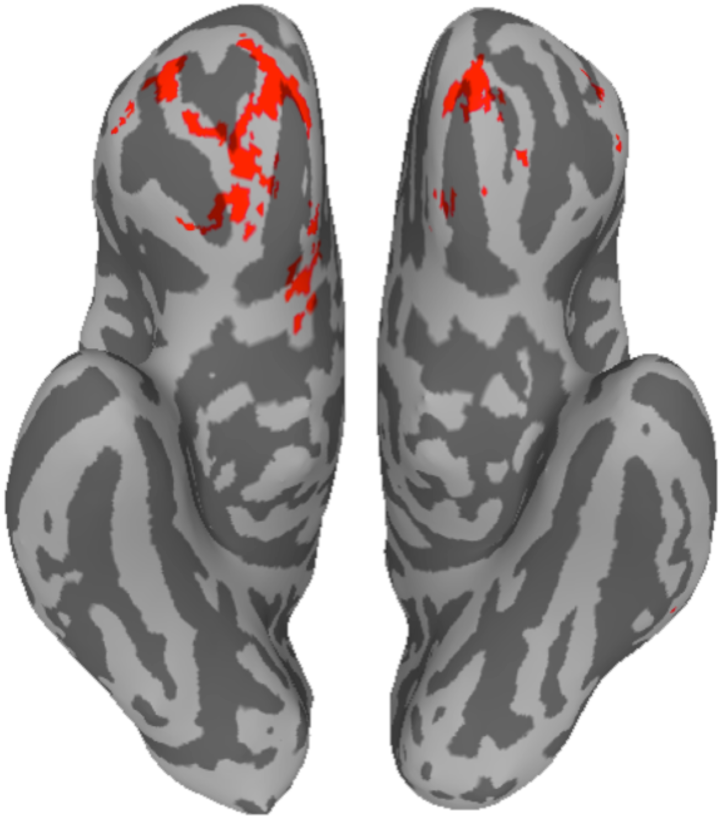
Wholebrain searchlight result. *Brain areas that passed both of the following criteria: (1) significantly higher representational similarity match with the logposterior model as compared with every other model from Table 3 except the posterior, the posterior from the feedbackRL model, and the entropy, at p < 0.05 with whole-brain correction for every comparison; (2) higher representational similarity match with the logposterior compared to the entropy, at p < 0.05 uncorrected. The map is displayed on the orbital/ventral surface of an inflated brain*.

## DISCUSSION

Because the underlying structure of the world is often not directly observable, we must make inferences about the underlying situations or “latent causes” that generate our observations. The statistically optimal way to do this is to use Bayes rule to infer the posterior distribution over latent causes. Based on previous studies implicating the orbitofrontal cortex (OFC) in the representation of the current context or situation (related to the ideas of “state” in studies of reinforcement learning, and “schemas” in studies of episodic memory), we hypothesized that the OFC might represent a posterior probability distribution over latent causes, computed using approximately Bayesian inference. To test this, we asked participants to make inferences about the probability of possible situations, in an environment where the situation probabilistically generated their observations.

Using representational similarity analysis of fMRI activity during the inference task, we found that patterns of activity in the suborbital sulcus within the OFC were indeed best explained as representing a posterior distribution over latent causes. Searchlight analyses implicated OFC more generally in this representation. Furthermore, participants’ behavioral performance showed that they had access to a full posterior distribution over the latent causes for their choices; using trial-by-trial model fitting, we showed that participants’ behavior was best explained as using Bayesian inference.

Our study provides evidence that the OFC represents a full posterior distribution over situations, as opposed to the best guess of the situation (the maximum a posteriori; MAP) or other summary measures of the distribution such as the overall uncertainty. We operationalized uncertainty as the entropy of the distribution—the highest entropy occurs when the distribution is completely flat (i.e., the participant is maximally uncertain about which latent cause generated the observations), and the lowest entropy occurs when the distribution is fully loaded on one latent cause (i.e., the participant is absolutely certain about which latent cause generated the observations). Our similarity analyses showed the entropy to have widespread positive similarity match in many areas of cortex, which we might expect because entropy should be correlated with the difficulty of the task, and so entropy might therefore be correlated with greater overall activity in many regions of the brain. Nonetheless, in greater than 95% of our bootstrap samples, activity in the OFC was better explained by the posterior distribution than by the entropy. Furthermore, searchlight analyses showed the specificity of this result.

Our results, using multivariate analysis, build on previous fMRI studies that have used univariate analyses in OFC to investigate a range of summary statistical quantities that are related to the posterior distribution, but which do not capture the full distribution. These studies have shown that univariate activation of the ventromedial PFC (which includes or is similar to our ROI) is correlated with a variety of summary statistics, e.g. expected reward (Ting et al, 2015), reward uncertainty (Tobler et al, 2007; Critchley et al, 2001), variance of the prior distribution in a sensory task (Vilares et al, 2012), and marginal likelihood of the current stimulus (d’Acremont et al 2013). Our experiment employed several key features — (a) multivariate neural analysis (b) four different latent causes, and (c) dissociation of latent cause from both reward and motor plan — that allowed us to identify orbitofrontal representation of a full posterior distribution over latent causes that was separate from value, and which explained neural activity in the area better than any single summary statistic that we tried. Our result may therefore explain why evidence for different summary statistics was found in different studies—these are all components of the full posterior distribution, or correlates of it.

Our study also builds on previous work in the fields of reinforcement learning and episodic memory that has implicated the OFC in representations of the current situation or context. In reinforcement learning, a belief distribution over states is necessary for optimal decision-making when the state of the world is not directly observable (partially observable Markov decision processes; Kaelbling et al, 1998). The OFC has long been implicated in reinforcement learning and decision-making in a wide range of settings; a recent review provides a unifying explanation for these results by postulating that the OFC represents inferred states in partially observable situations (Wilson et al, 2014). In theories of episodic memory, it is believed that we organize our memories according to an inferred “schema” that specifies the situation and stores previously learned relationships that a new memory can be incorporated into (Tse et al, 2007; Hupbach et al, 2008). These schemas seem to be represented or processed in the ventromedial prefrontal cortex (vmPFC, an area of the brain that is similar to our ROI; for reviews, see Schlichting and Preston, 2015; van Kesteren et al, 2012; Ranganath and Ritchey, 2012). For example, Tse et al (2011) showed evidence that activation of rat mPFC is highest immediately after memory encoding that should involve incorporating new information into existing schemas. Ezzyat and Davachi (2011) showed that greater activation of ventromedial PFC in humans during memory encoding is correlated with how strongly those memories are associated with other memories in the same “event”, consistent with the idea that vmPFC is involved in schemas that are bound to memories. Our results confirm the involvement of OFC in representations of the current situation, and additionally show that this representation in OFC takes the form of a *distribution* over possible situations.

Finally, our work also builds on previous studies investigating neural circuits involved in the “weather prediction” task, very similar to ours, in which one of two “weather” outcomes is probabilistically predicted by sequences of cards. Knowlton et al (1996) implicated the striatum in the learning of these probabilistic associations. In our task, participants learned the animal likelihoods outside the MR scanner, and thus we could not assess the brain areas involved in the learning phase. However, our results are compatible with Knowlton et al’s insofar as the OFC may use associations learned by the striatum (in our experiment, the animal likelihoods) to make inferences when presented with new observations (in our experiment, the “photographs” task). More recently, Yang and Shadlen (2007) used the weather-prediction task to show representation of a decision variable in parietal cortex that took the form of the log likelihood ratio between two options. In our experiment, we decorrelated the posterior probability from both decision variables and stimulus-reward associations, and we also investigated representations of the posterior probability over latent causes *before* the decision period. We conjecture that the OFC contains representations of the current state or situation in terms of a posterior distribution over the possible states, a representation that is likely used by downstream areas, e.g. parietal cortex, for decision making.

Previous work on the weather-prediction task also showed that most individuals employed heuristic strategies in inferring the weather (Gluck et al, 2002). In our experiment, we explored several heuristic models of participants’ inference, but were not able to find any that predicted participants’ behavior better than the optimal Bayesian models. There are several reasons why our task may have discouraged the use of heuristics. First, the animal likelihoods in our experiment were designed to avoid one-to-one mappings between observations and latent causes. Second, the task environment had four possible latent causes (instead of two), and the task itself required rank-ordering all four latent causes rather than just estimating the maximum a posteriori, thus increasing complexity and leading to the inadequacy of simple heuristics. Finally, we provided participants with a large amount of training on the probabilistic model of the world, so that heuristics may have been less necessary.

The posterior distribution we found in the OFC was best modeled as being represented in log space. Representation in log space may be advantageous because addition can then replace the multiplicative operation required to accumulate evidence in non-log space (e.g. across animal presentations, in our experiment); the ability of neurons to add is well-characterized, while it is less clear to what extent neurons can multiply (Yuste and Tank, 1996; Peña and Konishi, 2001; Gabbiani et al, 2002). Indeed, neural representation in log space is common in many domains, e.g. decision variables (Yang and Shadlen, 2010), time (Gibbon, 1977) and numbers (Longo and Lourenco, 2007).

To summarize, we designed a task in which participants’ observations were probabilistically generated by unobserved “situations” or “latent causes”, and found evidence that OFC represents a probability distribution over possible latent causes. A representation of the log posterior distribution explained OFC activity better than alternatives such as the best guess of the current situation, or overall uncertainty in the current situation. This finding was further supported by behavioral evidence that participants had access to the full probability distribution for decision-making, and used Bayesian inference to compute the probability distribution. Our results may explain why previous studies of OFC have found evidence for representation of various summary statistical quantities in OFC (these are in fact components of the full posterior probability distribution). Our results may also unify findings from disparate literatures on reinforcement learning and episodic memory, which separately implicate the OFC in representations of the current situation.

## ACKNOWLEDGEMENTS

This work was supported by National Science Foundation/National Institutes of Health Collaborative Research in Computational Neuroscience grant number NSF IIS-1009542, National Institutes of Health grant 2T32MH065214, and U.S. Army Research Office grant number W911NF1410101.

